# Highly efficient site-specific mutagenesis in Malaria mosquitoes using CRISPR

**DOI:** 10.1101/157867

**Authors:** Ming Li, Omar S. Akbari, Bradley J. White

## Abstract

*Anopheles* mosquitoes transmit at least 200 million annual malaria infections worldwide. Despite considerable genomic resources, mechanistic understanding of biological processes in *Anopheles* has been hampered by a lack of tools for reverse genetics. Here, we report successful application of the CRISPR/Cas9 system for highly efficient, site-specific mutagenesis in the diverse malaria vectors *Anopheles albimanus*, *Anopheles coluzzii*, and *Anopheles funestus*. When guide RNAs and Cas9 are injected at high concentration, germline mutations are common and usually bi-allelic allowing for the rapid creation of stable, mutant lines for reverse genetic analysis. Our protocol should enable researchers to dissect the molecular and cellular basis of anopheline traits critical to successful disease transmission, potentially exposing new targets for malaria control.

## INTRODUCTION

*Anopheles* mosquitoes are the exclusive vectors of mammalian malaria (White et al., 2011). Over the past decade, human malaria deaths have declined by nearly 50% primarily due to increased use of insecticides that target the mosquito vector (Bhatt et al., 2015). However, emerging physiological and behavioral resistance in *Anopheles* populations threatens the sustainability of insecticidal control (David et al., 2005; Edi et al., 2014; Ranson and Lissenden, 2016; Sougoufara et al., 2014). In order to maintain and extend the hard won progress of the past decade, novel vector control strategies need to be developed and combined with traditional chemical control. The development of new tools in the fight against malaria mosquitoes is contingent upon improved mechanistic knowledge of myriad mosquito biological processing including blood feeding, gametogenesis, gustation, immunity, olfaction, and metabolism, among many others.

In 2002, the African malaria mosquito *Anopheles gambiae* was the second arthropod to have its genome sequenced and, more recently, the genomes of 16 other anophelines were sequenced (Holt et al., 2002; Neafsey et al., 2013). Despite considerable genomic resources, progress in dissecting the molecular and cellular biology of malaria mosquitoes has been slow, primarily due to the difficulty in performing reverse genetic techniques that are routine in model organisms. Currently, the vast majority of *Anopheles* genes have no known function (Giraldo-Calderón et al., 2015), impeding the development of novel vector control strategies reliant upon understanding how individual genes contribute to the biology of the mosquito. Previously, genome editing in *Anopheles* relied on either transposon-based transgenesis with no control over where an insertion occurred (Carballar-Lejarazú et al., 2013; Grossman et al., 2001; Meredith et al., 2011; Nolan et al., 2002; Perera et al., 2002; Pondeville et al., 2014) or highly inefficient and expensive, site-specific genome editing technologies such as zinc finger nucleases or transcription activator-like effector nucleases (TALENs) (Smidler et al., 2013; Windbichler et al., 2007). Recently, the CRISPR/Cas9 genome editing technique has been successfully applied to a diversity of organisms (Barrangou and Doudna, 2016; Bassett et al., 2013; Basu et al., 2015; Dong et al., 2015; Gantz et al., 2015; Hall et al., 2015; Hammond et al., 2016; Hsu et al., 2014; Kistler et al., 2015; Li et al., 2017; Port et al., 2014; Sharma et al., 2017; Staahl et al., 2017). With this technology, researchers can directly edit or modulate DNA sequences, allowing them to study the function of genes *in vivo* (Hsu et al., 2014). When used for site directed mutagenesis, Cas9 protein and a small guide RNA (sgRNA) that is complementary to a target sequence in the genome are delivered to germ cells. The Cas9 and sgRNA complex, bind to the target sequence, and cause a double strand break, which will be repaired through non-homologous end joining (NHEJ) resulting in mismatches and indels relative to wild type sequence (Basu et al., 2015; Maruyama et al., 2015). When exons are targeted, such mutagenesis will often result in premature stop codons or frame shifts that disrupt protein function. Despite high mutagenesis efficiency in other organisms, it is unclear if the CRISPR/Cas9 system will prove to be efficient in *Anopheles* as egg injection alone often results in extremely high mortality and low transformation efficiencies, perhaps due to the inherent fragility of the eggs themselves. Here, we report successful development of an efficient site-specific mutagenesis protocol using the CRISPR/Cas9 system in various anophelines, facilitating reverse genetics in this important group of disease vectors.

## RESULTS

In order to rapidly and easily detect successful CRISPR/Cas9 mutagenesis, we wanted to target a gene where knockout of only a single allele produces a visible phenotype. However, no dominant visible mutations for *Anopheles* have been previously reported. Thus, we chose to target the *white* gene, which codes a protein critical for eye pigment transport (Besansky et al., 1995). Knockout of the white gene results in a change from wild type red eye color to white (unpigmented) eye color – a simple phenotype to score. Although the white gene is recessive, it is located on the X chromosome and thus hemizygous in male anophelines (XY sex determination system), meaning that successful knockout of a single allele in males will result in the white-eye phenotype.

### Mutagenesis efficiency is concentration and sgRNA dependent

*Anopheles coluzzii* belongs to the *Anopheles gambiae* complex, which includes a number of major African malaria vectors. To determine the efficacy of CRISPR/Cas9 mutagenesis in this species complex, we designed two sgRNAs targeting exon 2 of the *white protein* gene (ACOM037804). First, we used AcsgRNA1 to test how different concentrations of both the sgRNA and Cas9 protein effected mutagenesis rates. We found that both embryo survival and mutagenesis rate were sgRNA and Cas9 concentration dependent (Table 1). Greater than 50% of embryos survived control injections with only water, however, survival rates for embryos (37%) injected with even the lowest concentration of sgRNA and Cas9 decreased relative to control. With increasing concentrations of sgRNA and Cas9 embryo survival further decreased. Indeed, only 11% of embryos survived injections with the highest concentrations tested. Conversely, concentrations of sgRNA and Cas9 were positively correlated with mutagenesis rates; 46% of males injected with the lowest concentration had mosaic white eyes, while a remarkable 100% of males injected with the highest concentration had mosaic eyes (Figure 1, Table 1). Importantly, at higher injection concentrations, a majority of injected *females* also had mosaic eyes. Since the white gene is recessive, the production of mosaic females demonstrates that the CRISPR/Cas9 system can mutate both copies of diploid *Anopheles* genes. Notably, we also observed G0 injected males and females with completely white eyes, suggesting that vast majority of cells in the eyes were mutated. Based on the above results, we used an sgRNA concentration of 120ng/ul and a Cas9 protein concentration of 300 ng/ul, which balances survival and mutagenesis efficiency, to further explore the CRISPR/Cas9 system in *Anopheles*. To determine if sgRNA sequence has an effect on mutagenesis rate we compared AcsgRNA1 from above against a second sgRNA (AcsgRNA2) targeting white. We found that AcsgRNA1 (93%, 87%) produced mosaic G0 males and females at a much higher frequency than AcsgRNA2 (32%, 25%) suggesting that sgRNA sequence can have a large impact on mutagenesis efficiency (Table 2).

**Table 1.** Effect of sgRNA and Cas9 concentration on *An. coluzzii* survival and mutagenesis

| AcsgrNA1 | Cas9 | # Inj. | Survivors |  |  | Mosaic (%) |  |
| --- | --- | --- | --- | --- | --- | --- | --- |
|  |  |  | *M* | *F* | *Total (%)* | M (%) | F (%) |
| No injection | No injection | 300 | 137 | 118 | 255 (85) | 0 | 0 |
| Water | Water | 217 | 69 | 52 | 121 (56) | 0 | 0 |
| 30 (ng/ul) | 100 (ng/ul) | 185 | 31 | 38 | 69 (37) | 32 (46) | 0 |
| 60 | 200 | 251 | 48 | 33 | 81 (32) | 48 (59) | 0 |
| 120 | 300 | 219 | 31 | 16 | 47 (21) | 29 (94) | 12 (75) |
| 240 | 400 | 177 | 22 | 11 | 33 (19) | 20 (91) | 9 (82) |
| 480 | 500 | 228 | 12 | 12 | 24 (11) | 12 (100) | 10 (83) |

**Table 2.** G_0_ and G_1_ mutagenesis rates in three different *Anopheles* species.

|  | sgRNA | # Inj. | Survivors |  |  | Mosaic (%) |  | G<sub>0</sub> M x White F |  | White M x G<sub>0</sub> F |  | G<sub>0</sub> M x G<sub>0</sub> F |  |
| --- | --- | --- | --- | --- | --- | --- | --- | --- | --- | --- | --- | --- | --- |
|  |  |  | M | F | Total (%) | M (%) | F (%) | G<sub>1</sub> Mutant M (%) | G<sub>1</sub> Mutant F (%) | G<sub>1</sub> Mutant M (%) | G<sub>1</sub> Mutant F (%) | G<sub>1</sub> Mutant M (%) | G<sub>1</sub> Mutant F (%) |
| *Anopheles coluzzii* | AcsgrNA1 | 612 | 76 | 62 | 138 (23) | 71 (93) | 54 (87) | 991 (93) | 117 (91) | 851 (91) | 1232 (94) | 881 (86) | 939 (81) |
|  | AcsgrNA2 | 447 | 53 | 36 | 89 (20) | 17 (32) | 9 (25) | 1038 (88) | 1273 (84) | 751 (89) | 882 (91) | 751 (45) | 846 (49) |
| *Anopheles albimanus* | AasgrNA1 | 573 | 81 | 58 | 139 (24) | 74 (91) | 43 (74) | N/A |  | N/A |  | 1317 (60) | 1577 (62) |
|  | AasgrNA2 | 511 | 79 | 68 | 147 (29) | 0 | 0 | N/A |  | N/A |  | N/A | N/A |
| *Anopheles funestus* | AfsgRNA1 | 237 | 15 | 11 | 26 (11) | 8 (53) | 7 (64) | N/A |  | N/A |  | 53 (65) | 92 (71) |
|  | AfsgRNA2 | 352 | 21 | 16 | 37 (11) | 5 (24) | 4 (25) | N/A |  | N/A |  | 37 (51) | 62 (61) |

**Figure 1.**
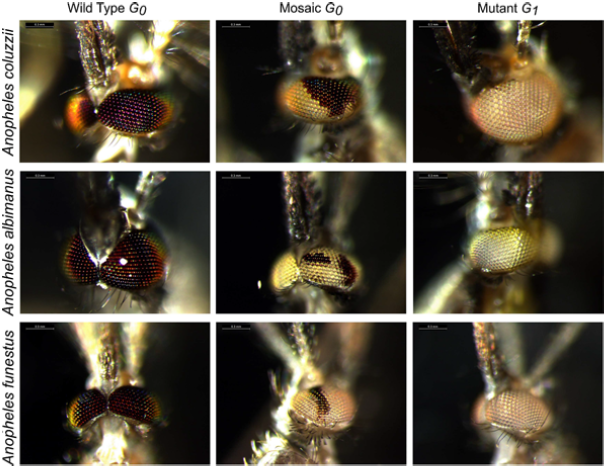
CRISPR/Cas9 efficiently generates heritable, site-specific mutations in diverse *Anopheles* mosquitoes. On the left, representative images of wild type anopheline eyes are shown for each species. In the center are representative G_0_ mosaic white-eyed mutant mosquitoes that were injected with sgRNA and Cas9 as embryos. On the right are representative homozygous white-eyed mutant G_1_ mosquitoes generated by crossing mosaic G_0_ male and female mosquitoes.

### Confirmation of Germline Mutations and Site-Specificity

While mosaic G0 mosquitoes can be used for reverse genetics, the creation of stable, mutant lines permits more thorough investigation of gene function. Thus, we wanted to determine the proportion of G0 mosaic-eyed *An. coluzzi* that possessed germline mutations. To obtain the germline mutation rate, we crossed G0 mosaic eyed males with females of an existing white-eye mutant line of *An. coluzzii* (M2) that was established more than 20 years ago (Benedict et al., 1996; Besansky et al., 1995; Mason, 1967). Hemizygous male progeny of this cross will all have white eyes since they inherit a maternal mutated white gene, however, homozygous females will only have white eyes if they inherit a mutant allele from both parents. A remarkable 93% of G0 mosaic males injected with AcsgRNA1 produced G1 females with white eyes, while 88% of G0 AcsgRNA2 mosaic males passed on white eye mutations to G1 female progeny (Table 2). To determine if female mosquitoes with mosaic eyes could also pass on the mutation, we performed a bulk cross of mosaic G0 males with mosaic G0 females and found that 83% (male 86%, female 81%) of the G1 progeny from AcsgRNA1 injected mosquitoes had fully white eyes, while 47% (male 45%, female 49%) of G1 progeny from AcsgRNA2 injected mosquitoes had fully white eyes (Table 2). We have now maintained multiple, white-eyed mutant lines in the laboratory for more than 15 generations, proving that the mutations introduced by CRISPR/Cas9 are highly stable. The combination of good G0 survival, biallelic mutation, and high germline transmission allows for the rapid creation of knockout *Anopheles coluzzii* lines using CRISPR/Cas9.

To confirm that the mosaic eye phenotype was caused by loss of function of the *white* gene, we performed T7 endonuclease I (T7EI) assay on five randomly chosen G0 AcsgRNA1 and AcsgRNA2 male mosquitoes with mosaic eyes. In the T7EI assay, T7 endonuclease will cut when NHEJ of the CRISPR-induced double strand break introduces a SNP or indel relative to the wild type allele, whereas no digestion will occur in mosquitoes with two wild type alleles. As expected, PCR fragments of the *white* gene from mosaic eye males were consistently cut into small bands by T7 endonuclease, while no activity was observed in non-mosaic male mosquitoes (Figure S2A-E), confirming that the white-eye phenotype is caused by disruption of *white* coding sequence. To sample the spectrum of mutations introduced by NHEJ, we performed Sanger sequencing of PCR products containing the two sgRNA target sites in G1 mosquitoes with white eyes. The sequencing results confirmed the presence of indels (Figure S1A-B) in all mutant mosquitoes that ranged in size from 2 to 54 base pairs. Finally, we screened for off-target activity of both sgRNAs by T7EI assay. Across three potential off target loci for both sgRNAs, no evidence of mutagenesis was detected in G0 mosaic males indicating high specificity of the sgRNAs (Figure S3).

### CRISPR/CAS9 activity in diverse Anopheles

To determine the applicability of the CRISPR/Cas9 system to diverse *Anopheles* species, we performed injections targeting *white* in *Anopheles albimanus* (a minor vector of malaria on South America) and *Anopheles funestus* (a major, understudied malaria vector in Africa) (Neasfey et al 2013). For each species, we designed two sgRNAs targeting *white* and injected the individual sgRNA (120 ng/ul) and Cas9 (300 ng/ul) directly into eggs.

As found in other studies, *Anopheles albimanus* survived injections at a rate comparable to *Anopheles coluzzii (Perera et al., 2002)*. For AasgRNA1, we found that 91% of G0 males had mosaic white eyes, while 74% of G0 females were mosaics. Interestingly, injection of AasgRNA2 produced no mosquitoes with mosaic eyes, further reinforcing the impact of sgRNA choice on mutagenesis efficiency. Since no previously generated white-eyed line of *An. albimanus* was available, we bulk crossed AasgRNA1 G0 mosaic males and females to determine germline mutation rates (Table 2). Over 61% of the G1 progeny from the cross possessed fully white eyes, suggesting high germline mutagenesis efficiency in *An. albimanus*. As with *An. coluzzii*, T7EI assays consistently detected mutations in white in G0 mosaic males (Figure S2) and sequencing showed indels in mosaic males ranging from 2 to 11 base pairs (Figure S1). Additionally, no mutations were detected using T7EI assays at three potential off target sites for AasgRNA1 (Figure S3).

Survival rate of *Anopheles funestus* embryos injected with either of two sgRNAs targeting the white gene (AfsgRNA1 and AfsgRNA2) was less than half that of *An. coluzzii* and *An. albimanus*. (Table 2) We attribute the lower survival rate to the unique morphology of *An. funestus* eggs at the poles (Figure 2), which makes injection challenging. Despite lower survival, a high proportion of G0 males (53% and 24%) and females (64% and 25%) for both sgRNAs displayed mosaic white eyes. Bulk crossing of G0 male and female mosaics produced 67% (AfsgRNA1) or 57% (AfsgRNA2) G1 progeny with fully white eyes demonstrating high germline mutagenesis rates. As with previous species, the T7EI assay consistently identified mosaic males (Figure S2) and Sanger sequencing revealed diverse indels (2 to 34 base pairs) in mutated males (Figure S1). No mutagenic activity was detected for either sgRNA at the three most likely off target genomic sites (Figure S3).

**Figure 2.**
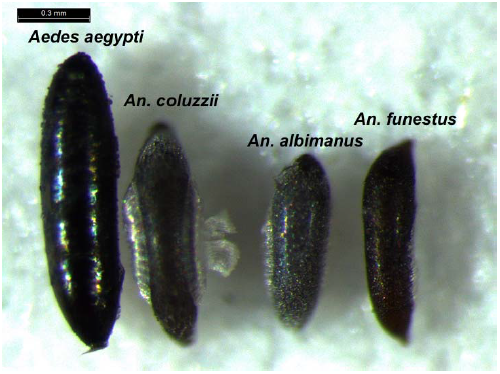
Morphology of eggs differs dramatically among anophelines. Eggs of the three species of *Anopheles* used in this study alongside an egg of the yellow fever mosquito *Aedes aegypti* for size comparison. Note the difference in pole shape between *Anopheles albimanus* and *Anopheles funestus* eggs, which likely contributes to differences in both survival and mutagenesis rates between these two species.

## DISCUSSION

The CRISPR/Cas9 system offers the possibility of precise, efficient, and cost effective mutagenesis in non-model organisms (Barrangou and Doudna, 2016). While considerable genomic resources have been developed for malaria mosquitoes, no efficient tools for performing reverse genetics in these species exist, slowing the development of genetically based vector control. Studies of the CRISPR/Cas9 system in the distantly related (diverged 145-200 million years ago) mosquito *Aedes aegypti* have demonstrated G0 mutagenesis rates between 3 – 50% with high variability among injection operators and different sgRNAs (Basu et al., 2015; Kistler et al., 2015). While no systematic studies of CRISPR/Cas9 mutagenesis rates on any anopheline mosquitoes have been conducted, a few groups developing gene drive related technologies (Champer et al., 2016) have recently reported high rates of mutagenesis when guide RNAs and Cas9 were directly integrated into the genome of two *Anopheles* species (Galizi et al., 2016; Gantz et al., 2015; Hammond et al., 2016).

In summary, we report remarkably high rates of survival and mutagenesis in three different *Anopheles* species co-injected with Cas9 protein and sgRNAs targeting the *white* gene. Importantly, we describe the first, successful genetic engineering of *An. funestus*, demonstrating that the CRISPR/Cas9 system may even be useful in species where previous genome editing techniques proved too inefficient for practical use. Additionally, since high concentrations of sgRNA and Cas9 result in biallelic mutations, stable mutant lines can be rapidly generated even when no visible marker is present. The following procedure can be used to generate such lines:1) inject sgRNA and Cas9 into mixed sex eggs, 2) cross G0 survivors en masse, 3) isolate G0 females into individual ovicups, 4) screen 5 G1 larvae from each family for mutations (sequencing, T7EI, or an alternative assay), 5) conduct full sibling mating of families in which all G1 larvae are mutants, and 6) confirm stable generation of a mutant line by sequencing of G2 mosquitoes. The ability to rapidly and consistently create stable knockout lines should greatly accelerate mechanistic research into key cellular and molecular pathways in malaria mosquitoes.

We note that cleavage efficiency of the sgRNA/Cas9 complex is target site dependent. In mammalian systems, it has been reported that the chromatin environment around the target site and certain features of the sgRNA sequence are major factors affecting the efficiency of DSB generation (Doench et al., 2014; Kuscu et al., 2014; Wang et al., 2015). Due to the limited number of sgRNAs we tested, we are unable to confirm whether these observations can be extended into *Anopheles*. However, the complete failure of AasgRNA2 to cause knockout is likely due to low thermodynamic stability of the sgRNA/Cas9 complex or secondary structure at the target site preventing binding (Bassett and Liu, 2014; Moreno-Mateos et al., 2015).

Having demonstrated the utility of the CRISPR/Cas9 system for site-specific mutagenesis of *Anopheles*, a logical next step is to systematically determine the efficiency of the system for integrating various sized constructs into anopheline genomes via HDR. The ability to conduct efficient deletion and addition of known sequences at specific genomic positions will greatly speed progress towards genetic methods, such as gene drive, for control of malaria vectors.

## Experimental Procedures

### Mosquito Strains

Four mosquito colonies were used in this study: *Anopheles coluzzii* wild-type strain NGS; *An. gambiae* white eyed mutant strain M2 (MRA-105); *An. albimanus* wild type strain *STECLA* (MRA-126) and *An. funestus* wild-type strain *FUMOZ* (MRA-127). Strains with accession numbers were obtained from the Malaria Research and Reference Reagent Resource Center (MR4). Mosquitoes were maintained in insectaries at the University of California, Riverside under standard conditions (White et al., 2013).

### sgRNA Design and Generation

Guide RNAs were designed by searching both the sense and antisense strand of exon 2 of the *white* gene (AGAP000553, AALB006905, and AFUN003538) for the presence of protospacer-adjacent motifs (PAMs) with the sequence NGG using ZIFIT (http://zifit.partners.org/ZiFiT/ChoiceMenu.aspx) and CRISPR Design (http://crispr.mit.edu/)(Xie et al., 2014). Linear, double-stranded DNA templates for sgRNAs were generated by performing template-free PCR with Q5 high-fidelity DNA polymerase (NEB). PCR conditions included an initial denaturation step of 98°C for 30 s, followed by 35 cycles of 98°C for 10 s, 58°C for 10 s, and 72°C for 10 s, following by a final extension at 72°C for 2 min. PCR products were purified with magnetic beads using standard protocols. Guide RNAs were generated by in vitro transcription (AM1334, Life Technologies) using 300 ng purified DNA as template in an overnight reaction incubated at 37°C. MegaClear columns (AM1908, Life Technologies) were used to purify sgRNAs, which were then diluted to 1 ug/ul, aliquoted, and stored at 80°C until use. All primer sequences are listed in Table S1. Recombinant Cas9 protein from *Streptococcus pyogenes* was purchased from PNA Bio (CP01) and diluted to 1 ug/ul in nuclease-free water with 20% glycerol and stored in aliquots at −80°C.

### Microinjection

Mixed sex pupae were allowed to eclose into a single (insert size) cage. After allowing five days for mating, females were offered a bovine bloodmeal using the Hemotek (model# PS5) blood feeding system. A minimum of 60 hours was allowed or oogenesis, after which ovicups filled with ddH20 and lined with filter paper were introduced into cages and females were allowed to oviposit in the dark. After ~15 minutes, the ovicup was removed and unmelanized eggs were transferred onto a glass slide and rapidly aligned against a wet piece of filter paper.Aluminosilicate needles pulled on a Sutter P-1000 needle puller and beveled using a Sutter BV-10 beveler were used for injections. An Eppendorf Femotojet was used to power injections, which were performed under a compound microscope at 100x magnification. Since eggs were injected prior to melanization, only 10-20 eggs were injected at a time, after which fresh eggs were obtained. After injection eggs were floated in ddH20 and allowed to hatch spontaneously.

### Mutation Screens

The white-eye phenotype of G0 and G1 mosquitoes was assessed and photographed under a Leica M165 FC stereomicroscope. To molecularly characterize CRISPR/Cas9 induced mutations, genomic DNA was extracted from a single mosquito with the DNeasy blood & tissue kit (QIAGEN) and target loci were amplified by PCR. For T7EI assays, 1 ul of T7 endonuclease (NEB) were added to 19 ul of PCR product and digested for 15 minutes at 37□ and visualized on a 2% agarose gel electrophoresis stained with ethidium bromide. To characterize mutations introduced during NHEJ, PCR products containing the sgRNA target site were amplified cloned into TOPO TA vectors (Life Technologies), purified, and Sanger sequenced at the UCR Genomics core.

## AUTHOR CONTRIBUTIONS

Conceived, designed, and performed experiments: ML and BJW. Analyzed data and wrote the paper: ML, OSA, BJW.

## ACKNOWLEDGMENTS

Funding was provided by NIH grants 1R01AI113248 and 1R21AI115271 to BJW. We thank Timothy Lo for help with injections and MR4, part of the BEI Resources Repository, for providing mosquito eggs to start colonies.

## SUPPLEMENTARY INFORMATION

**Figure S1.**
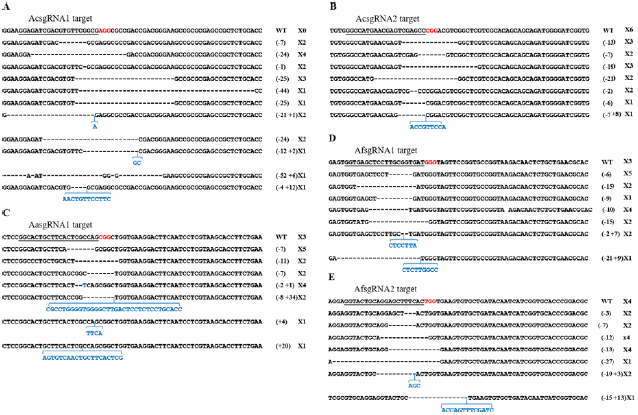
Repair of CRISPR-induced double strand breaks results in a variety of indels. Sequencing of cloned PCR products from G_0_ injected mosquitoes possessing mosaic eyes revealed a variety of insertions and deletions adjacent to the guide target site. For each sgRNA, top line represents WT sequence; PAM sequences (NGG) are indicated in red, and gene disruptions resulting from insertions/deletions are indicated in blue/dash.

**Figure S2.**
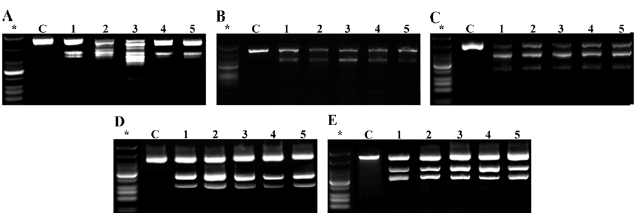
The T7 Endonuclease assay can be used for rapid detection of CRISPR-generated mutant alleles. For each successful guide RNA (A, AcsgRNA1; B, AcsgRNA2; C, AasgRNA1; D, AfsgRNA1; E, AfsgRNA2), PCR products from non-mosaic (C) and mosaic (1-5) mosquitoes were digested with T7 endonuclease. In all mosaic mosquitoes, partial digestion of the PCR product is evident, while in non-mosaic mosquitoes no digestion is visible.

**Figure S3.**
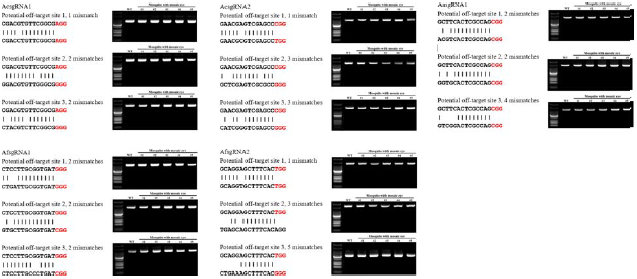
No evidence for off-target mutagenesis of sgRNAs. Three potential off target sites for each sgRNA were screened for mutagenesis activity by T7 endonuclease assay. PAM distal region sequence alignment of target locus and potential off-target loci. The potential offtarget sites of sgRNAs in different *Anopheles* mosquitoes are indicated, and the PAM sites are labeled in red. T7 Endonuclease I (T7E1) assay of potential off-target loci. “WT” represented wild type mosquito, number from 1 to 5 indicated 5 different mosquitos with mosaic eye phenotype. No digestion is visible in any of the lanes.

**Table S1.**
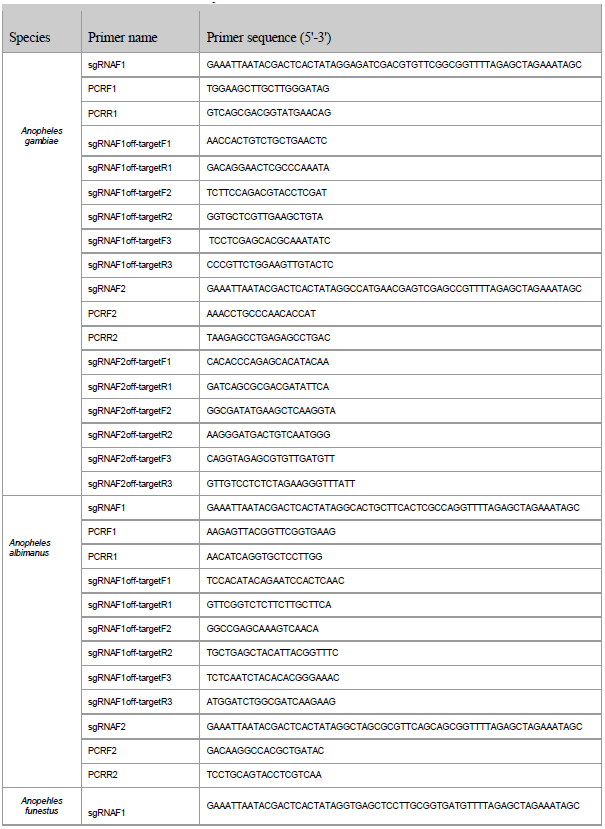
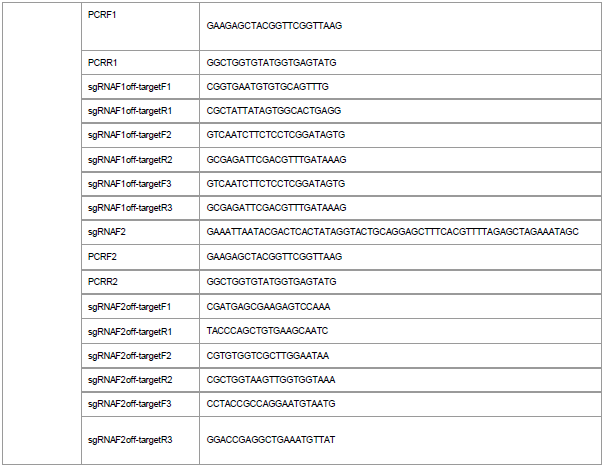
Primers used in this study.

